# Striatal activity reflects cortical activity patterns

**DOI:** 10.1101/703710

**Authors:** Andrew J Peters, Nicholas A Steinmetz, Kenneth D Harris, Matteo Carandini

## Abstract

The dorsal striatum is organized into domains that drive characteristic behaviors^1–7^, and receive inputs from different parts of the cortex^8,9^ which modulate similar behaviors^10–12^. Striatal responses to cortical inputs, however, can be affected by changes in connection strength^13–15^, local striatal circuitry^16,17^, and thalamic inputs^18,19^. Therefore, it is unclear whether the pattern of activity across striatal domains mirrors that across the cortex^20–23^ or differs from it^24–28^. Here we use simultaneous large-scale recordings in the cortex and the striatum to show that striatal activity can be accurately predicted by spatiotemporal activity patterns in the cortex. The relationship between activity in the cortex and the striatum was spatially consistent with corticostriatal anatomy, and temporally consistent with a feedforward drive. Each striatal domain exhibited specific sensorimotor responses that predictably followed activity in the associated cortical regions, and the corticostriatal relationship remained unvaried during passive states or performance of a task probing visually guided behavior. However, the task’s visual stimuli and corresponding behavioral responses evoked relatively more activity in the striatum than in associated cortical regions. This increased striatal activity involved an additive offset in firing rate, which was independent of task engagement but only present in animals that had learned the task. Thus, striatal activity largely reflects patterns of cortical activity, deviating from them in a simple additive fashion for learned stimuli or actions.

---

The cortex and the dorsal striatum are reciprocally connected, but the relationship between activity in the two structures is unclear, and so is the degree to which this relationship depends on learning. On the one hand, the cortex and the striatum can show correlated activity^20,29^ and undergo similar learning-related changes^23^. On the other hand, cortical and striatal activity may be differently affected by behavioral context^28^, becoming either more correlated^21,22^ or more dissimilar^30^ after learning. Different types of striatal response may depend differently on cortical input, possibly being entirely independent from the cortex^24,29^ or driven only by cortical cells related to specific stimuli^13^ or behaviors^22^, and only after learning^31^. Compounding this diversity of views is the fact that recordings have been performed piecewise: they could not parcel the striatum into functional domains and could not relate activity across the striatum to the activity of large regions of cortex. Yet, a comprehensive view of how cortical and striatal activity is related is a crucial step toward understanding how these two structures cooperate to drive behavior.

We investigated activity across associated cortical and striatal regions while mice executed learned visually-guided behavior. Mice were trained to orient towards visual stimuli: gratings of varying contrast presented on the left or right, which could be brought to the center by turning a wheel^32^ (**Fig. 1a, Extended Data Fig. 1**). While mice performed this task, we recorded activity simultaneously in the cortex and the striatum (n = 36 recordings across 5 mice, **Fig. 1b**). We measured cortical activity through widefield calcium imaging of GCaMP6s expressed transgenically in all excitatory neurons^33^ (**Fig. 1b top**). Fluorescence was corrected for hemodynamics, deconvolved, and aligned across animals (**Extended Data Fig. 2**). At the same time, we recorded activity across the dorsal striatum with a Neuropixels probe^34^ inserted diagonally from dorsomedial to dorsolateral striatum in the left hemisphere to target striatal domains putatively related to both vision and movement (**Fig. 1b bottom**). We sorted units with Kilosort2 (ref. ^35^) and estimated striatal boundaries from electrophysiological landmarks (**Extended Data Fig. 3**).

**Figure 1.**
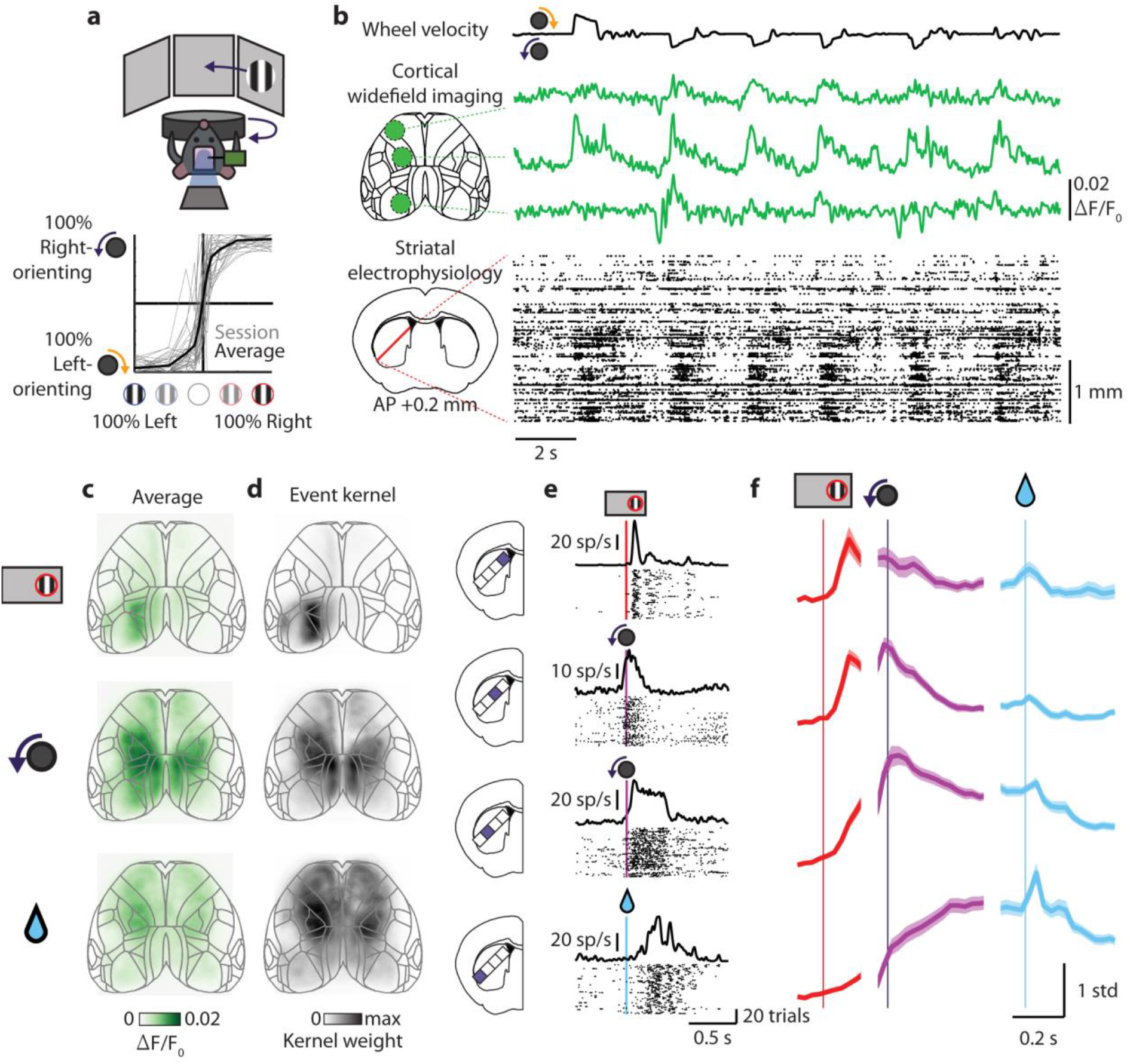
Cortex and striatum show spatial gradients of sensorimotor activity during visually guided behavior. **a**, Top, schematic of task and recording; bottom, task performance psychometric. **b**, Example recording session. Green traces: fluorescence from 3 ROIs; rasters: spikes recorded in the striatum plotted by depth. **c**, Trial-averaged cortical fluorescence 80 ms after stimulus onset for trials with 100% contrast right-hand stimuli (top), 0 ms after correct rightward-orienting movements (middle), and 80 ms after rewards (bottom), all during trials with reaction times less than 500 ms (from **Movie 1**). **d**, Frames from spatiotemporal kernels obtained by from cortical fluorescence on the same three task events (from **Movies 2-4**). **e**, Example units across the striatum aligned to contralateral (right-hand) stimuli (top), contralaterally (rightward)-orienting movements (middle two panels), and reward (bottom). f. Striatal multiunit activity grouped by depth relative to the striatum-piriform cortex border across experiments and aligned to contralateral stimuli (left), contralaterally-orienting movements (middle), and rewards (right), all during trials with reaction times less than 500 ms. Error bars represent SEM across recordings.

The spatial pattern of cortical activity evolved over the course of a trial, following the progression of sensory and motor events (**Fig. 1c-d, Movie 1**). Stimulus onset was followed by activity in visual cortex and medial frontal cortex contralateral to the stimulus (**Fig. 1c, top**); movement onset was accompanied by bilateral activity in retrosplenial cortex and limb somatomotor cortex (**Fig. 1c, middle**), and reward onset was followed by bilateral activity in the orofacial somatomotor cortex, likely due to licking and mouth movements (**Fig. 1c, bottom**). To isolate the activity related to each event type, we used regression analysis to model widefield fluorescence as a sum of spatiotemporal kernels triggered on each task event (visual stimulus onset, movement onset, go cue, and outcome events). The spatial profiles of the kernels matched the activity patterns seen in average movies: visual stimulus responses were concentrated in contralateral visual cortex and more weakly in medial frontal cortex (**Fig. 1d, top, Extended Data Fig. 4a, top row, Movie 2**); movement responses were largely bilateral and strongest in retrosplenial, medial frontal, and limb somatomotor cortex (**Fig. 1d, middle, Extended Data Fig. 4a, middle row, Movie 3**); and reward responses were bilateral and strongest in the orofacial somatomotor cortex (**Fig. 1d, bottom, Extended Data Fig. 4a, bottom row, Movie 4**). Go cue responses were concentrated over parietal cortex (our imaging region did not include auditory cortex), but were present only on uncommon trials when mice waited for the go cue to begin turning the wheel (**Extended Data Fig. 5a, Movie 5**). Summed together, these kernels successfully fit the fluorescence measured across the cortex within single trials (**Extended Data Fig. 4b**).

Echoing the progression of sensorimotor activity observed in the cortex, the task elicited a progression of activity from dorsomedial to dorsolateral striatum (**Fig. 1e-f**). Specifically, we found stimulus-locked activity in the dorsomedial striatum, movement-locked activity in the dorsocentral striatum, and reward-locked activity in the dorsolateral striatum, both in single neurons (**Fig. 1e**) and in pooled multiunit activity (**Fig. 1f**).

Thus, during task performance, activity flowed from posterior to anterior cortex, at the same time as from medial to lateral striatum. We hypothesized that this simultaneous flow of activity across the cortex and the striatum mapped onto the pattern of corticostriatal connections known from anatomy^8,9^.

We characterized the functional relationship between activity in the cortex and the striatum, and found that it obeys a topographic arrangement (**Fig. 2a**). We used ridge regression to estimate a spatiotemporal kernel that (when convolved with cortical fluorescence over a ±500 ms window) optimally predicted multiunit activity in each striatal location. This approach is similar to spike-triggered averaging but limits the impact of correlations induced by the task design, such as the temporal overlap of visual and movement responses. The resulting spatial maps of preferred cortical activity showed clear and diverse topographic patterns across striatal locations (**Fig. 2a**).

**Figure 2.**
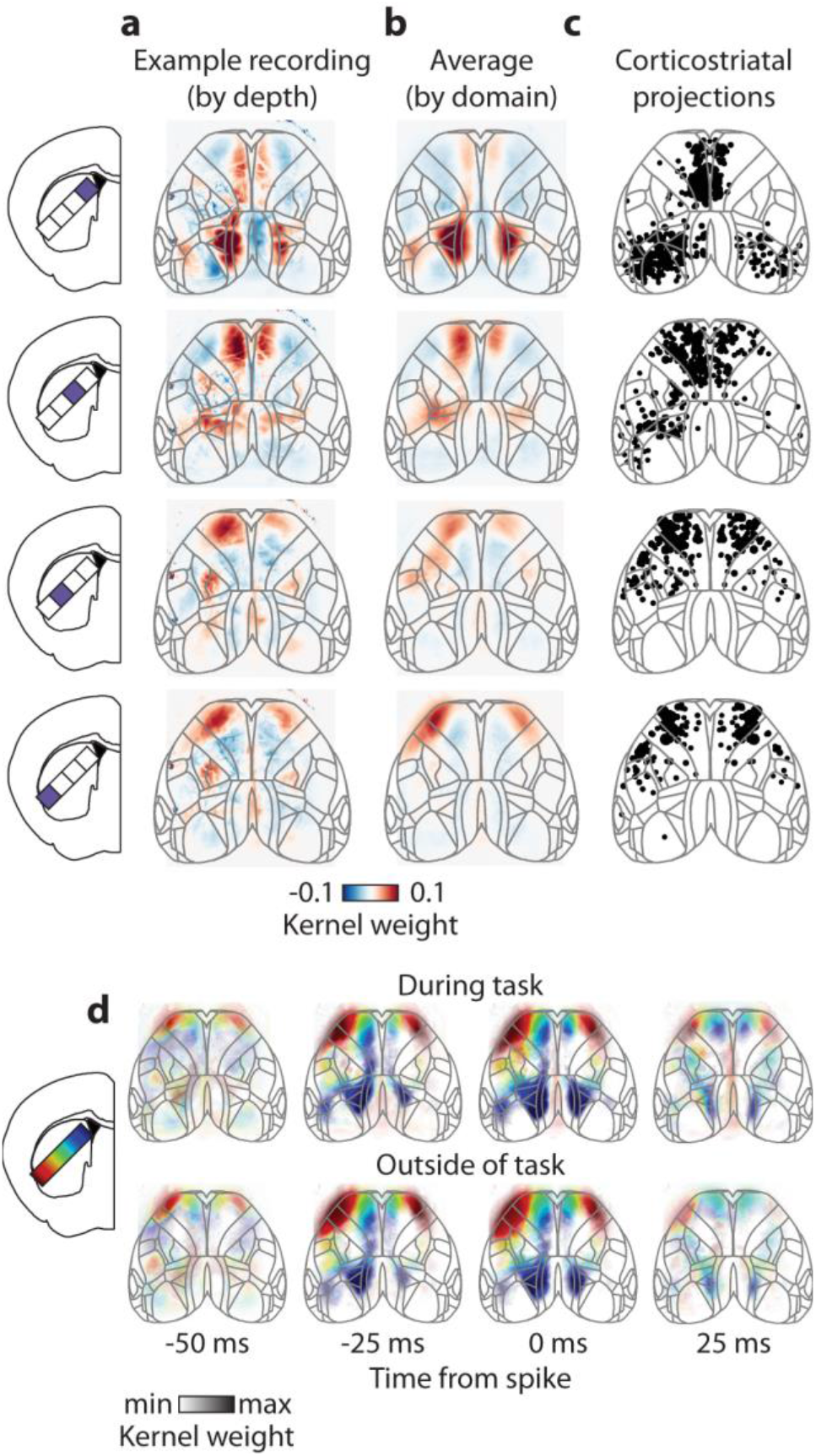
Striatal domains are topographically correlated with connected cortical regions. **a**, Example frames from spatiotemporal kernels of cortical activity optimally predicting striatal multiunit activity at four depths, in one recording session, time lag of 0 s shown as a subset of the full cortical kernel ranging from −500 ms to +500 ms relative to striatal firing. **b**, Similar analysis, averaging over each striatal domain across all recordings (from **Movie 6**). **c**, Anatomical organization of corticostriatal projections from the Allen connectivity database^35^, each dot represents a cortical injection site that yielded axons in each corresponding striatal location, dot size represents relative projection density, data taken from both hemispheres and oriented to indicate cortical projections to the left-hand striatum. **d**, Cortical activity kernels calculated from mice performing the task (top) or passively viewing sparse noise stimuli (bottom), shown across multiple time lags relative to striatal spikes. Colors represent center-of-mass striatal depth for each pixel weighted by kernel amplitude, with transparency normalized to the maximum amplitude across time.

The topographic relationship between cortical and striatal activity was stereotyped across experiments, allowing us to define striatal domains by their preferred cortical maps (**Fig. 2b, Extended Data Fig. 5, Movie 6**). We estimated preferred cortical maps for every 200 μm striatal segment in every recording, which produced a common set of maps across experiments (**Extended Data Fig. 6**). By clustering all of these maps into four groups, we could then assign each striatal segment into one of four striatal domains. This allowed us to estimate the borders between striatal domains within recordings and functionally align activity into common domains across recordings. We then found the average preferred cortical map for each domain across recordings, which showed that the mediolateral axis of the striatum was related to the caudorostral axis of the cortex, mirroring the progression of activity observed during the task (**Fig. 2b, Movie 6**).

The functional maps relating cortical and striatal activity matched the topographic pattern of anatomical projections from the cortex to the striatum and were contextually robust (**Fig. 2c-d**). To examine the pattern of anatomical corticostriatal projections to our four striatal domains we queried the Allen Mouse Brain Connectivity Atlas^36^ (**Fig. 2c**). The resulting anatomical maps resemble our functional maps (**Fig. 2b**), suggesting that corticostriatal projections likely provide the substrate for our observed functional associations. Consistent with feedforward connectivity from the cortex to the striatum, the temporal component of our preferred cortical patterns of activity slightly preceded striatal spiking and was restricted to about a 30 ms window (**Fig. 2d, top row, Movie 6**). Importantly, the spatiotemporal relationship between cortical and striatal activity was consistent independently from task engagement: the preferred cortical patterns were the same when constructed from activity during the task (**Fig. 2d, top row**) or when not engaged in the task (**Fig. 2d, bottom row**). Together, these results suggest that corticostriatal projections provide a robust channel by which spatial patterns of cortical activity drive activity in specific striatal domains, independently of behavioral context.

The progression of activity across the striatum during a trial reflected a gradient of sensorimotor correlates across striatal domains (**Fig. 3a-c, Extended Data Fig. 7–8**). To characterize striatal responses during the task, we used regression to obtain kernels relating task events to striatal activity (as in **Fig. 1d** for the cortex). The medial domain had contrast-dependent responses to contralateral stimuli (**Fig. 3a-b, row 1, Extended Data Fig. 7, row 1**), the centromedial domain had responses straddling movement onset and favoring contralaterally-orienting movements (**Fig. 3a-b, row 2, Extended Data Fig. 7, row 2**), the centrolateral domain had responses after movement onset favoring contralaterally-orienting movements (**Fig. 3a-b, row 3, Extended Data Fig. 7, row 3**), and the lateral domain had responses following reward (**Fig. 3a-b, row 4, Extended Data Fig. 7, row 4**). These task-related kernels together provided a faithful summary of activity on a trial-by-trial basis (**Fig. 3c, Extended Data Fig. 8a**), indicating that striatal activity represented a sum of visual (medial), motor (central), and reward (lateral) responses.

**Figure 3.**
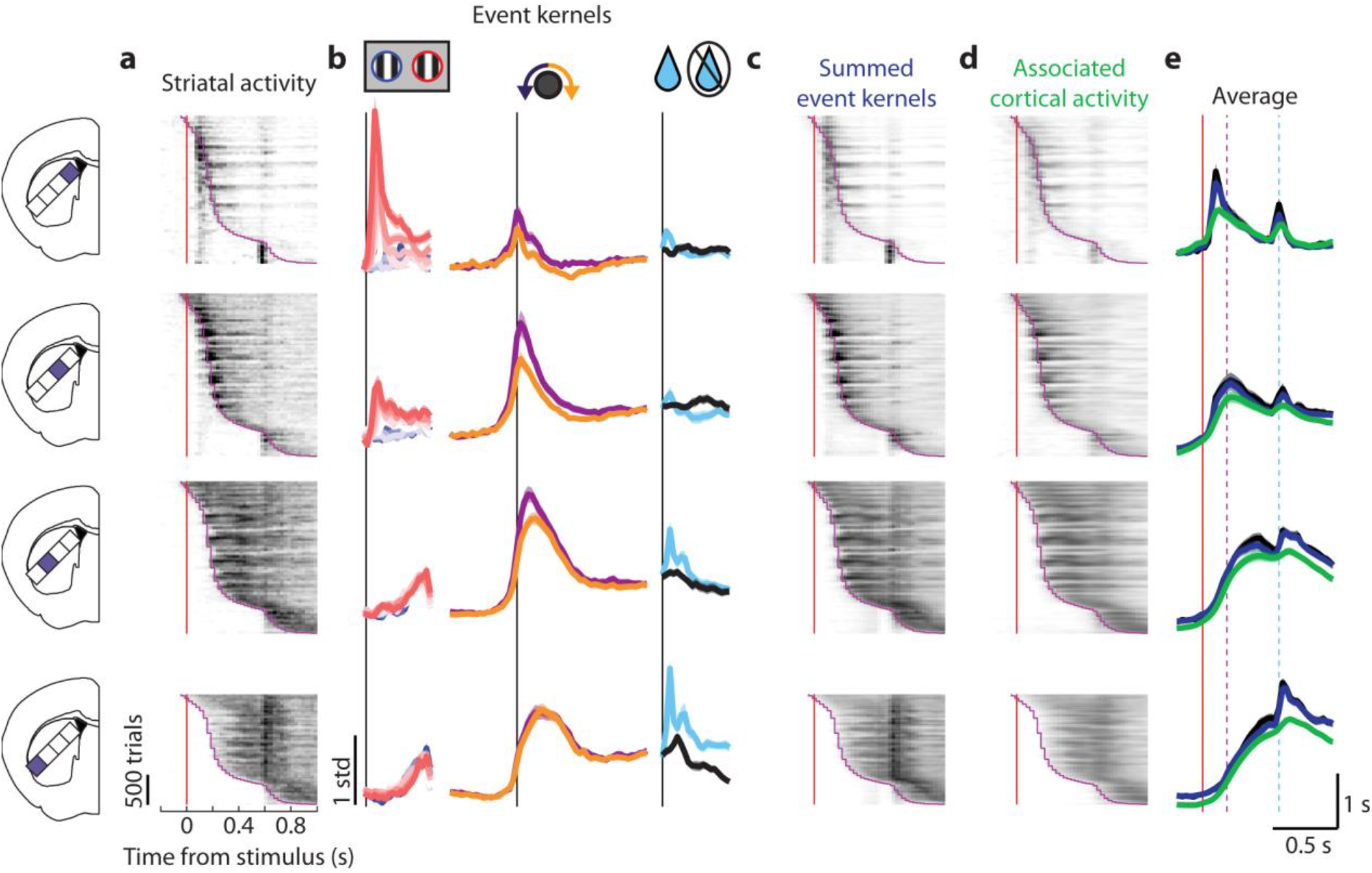
Striatal domains exhibit a sensorimotor gradient of activity consistent with both task responses and cortical activity. **a**, Activity for each striatal domain across all trials from all recordings with contralateral stimuli and contralaterally-orienting movements. Trials are sorted vertically by reaction time; red line: stimulus onset, purple curve: movement onset. Activity within each timepoint smoothed with a running average of 50 trials to display across-trial trends. **b**, Kernels from regressing task events to striatal domain activity for stimuli (left), movements (middle), and outcome (right). **c**, Prediction of activity in each striatal domain by summing kernels for task events. Trials ordered vertically as in (a). **d**, Prediction of striatal activity from cortical activity, displayed similarly to (a,c). **e**, Activity in each striatal domain averaged across trials (from a, black), compared to prediction from task events (from c, blue), and predicted from cortical activity (from d, green); red line: stimulus onset, purple line: average movement onset, cyan line: average reward time. Error bars represent SEM across recordings.

The diverse task-related activity of different striatal domains was closely predicted by the activity of their associated cortical regions (**Fig. 3d-e**). If the striatal domains reflect activity of their preferred cortical maps, it should be possible to predict their task correlates based on the task correlates of different cortical regions (shown in Fig. 1), together with the relationship between cortical and striatal activity (shown in Fig. 2). We computed the summed activity across cortical regions using the preferred spatiotemporal maps of cortical activity for each striatal domain (shown in Fig. 2b). This prediction was made solely from cortical activity, without using any explicit information about the task. We found that domain-associated cortical activity largely matched striatal activity on a trial-by-trial basis (**Fig. 3d-e, Extended Data Fig. 8a**). Moreover, we found that predictions of striatal activity were superior when they were made from full preferred cortical maps rather than from individual spots in cortex (**Extended Data Fig. 8a, green vs. orange lines**). These observations indicate that striatal task correlates are largely shared with the cortex and not further patterned by subcortical circuitry.

Nonetheless, specific striatal domains were more active for contralateral stimuli and movements than predicted from their associated regions of cortex (**Fig. 3e, Extended Data Fig. 8b**). For example, the dorsomedial striatum showed larger responses to contralateral stimuli (**Fig. 3e row 1, Extended Data Fig. 8b row 1 column 1**), and the dorsocentral striatum showed larger activity around contralaterally-orienting movements (**Fig. 3e row 2, Extended Data Fig. 8b row 2 column 2**). These deviations between striatal activity and cortical prediction suggest that there are event-specific divergences from an otherwise consistent corticostriatal relationship. In other words, while a simple readout of cortical activity can describe much of the activity within the striatum, specific events trigger an increase in striatal activity not explained by the normal corticostriatal relationship.

We tested the hypothesis that the corticostriatal relationship diverges around particular events by comparing trials with different task events but matched levels of cortical activity (**Fig. 4a**). This was made possible by the substantial variability in activity across trials. For example, the cortical region associated with the dorsomedial striatal domain comprised primarily higher visual cortex, and responded most strongly to contralateral visual stimuli. Nevertheless, trial-to-trial variability meant that activity in this cortical region on some trials with contralateral stimuli equaled activity on other trials with 0% contrast (invisible), or ipsilateral stimuli. We binned trials with matched cortical activity across stimulus conditions (**Fig. 4a, top row, green lines**), and plotted mean striatal activity for each combination of stimulus and cortical activity bin (**Fig. 4a, bottom row, gray lines**). If striatal activity were a fixed function of cortical activity, this analysis would yield an identical relationship between striatal and cortical activity, regardless of sensory or motor conditions.

**Figure 4.**
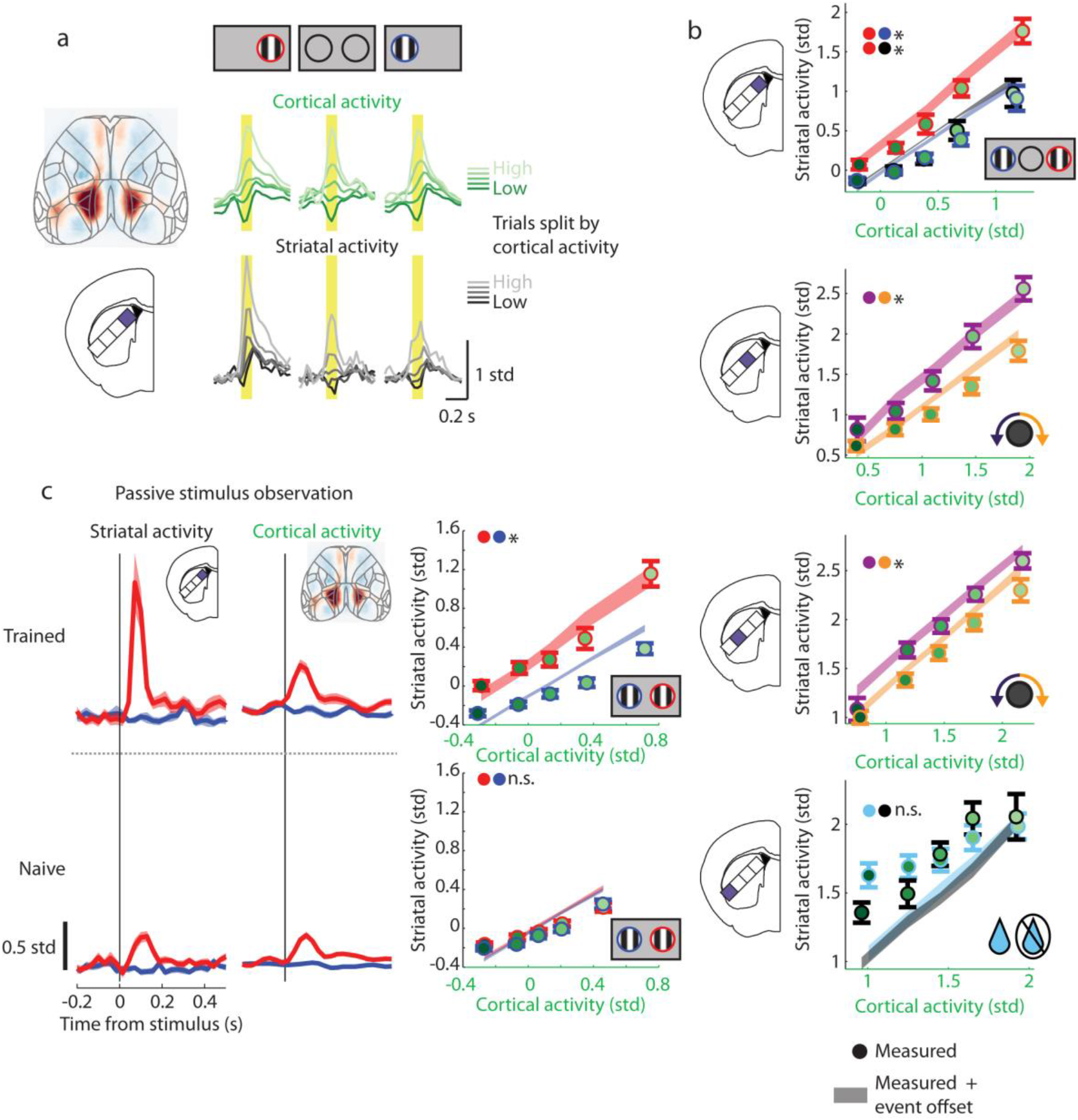
Striatal activity deviates from associated cortical activity for stimuli and movements in a learning-dependent manner. **a**, Top, cortical widefield activity was weighted by the spatiotemporal kernel corresponding to the dorsomedial striatal domain (left), and trials were divided into 5 bins across each stimulus type according to the amount of cortical activity following stimulus onset on that trial. The smallest and largest 5% of trials were excluded to minimize outlier effects. Different shades of green curves show cortical activity averaged over these 5 bins, for contralateral stimuli (left column), 0% contrast stimuli (center column), and ipsilateral stimuli (right column). Yellow bar shows time window used to assign trials to bins (50-150 ms following stimulus onset). The similar shape of these curves for each stimulus type indicates that binning has equalized cortical activity. Bottom, dorsomedial striatal activity, averaged over the same trials used to compute the green traces above. **b**, Activity in each striatal domain (y-axes) as a function of associated cortical activity (x-axes, corresponding to the 5 cortical activity bins), and task events relevant to each striatal domain (indicated by different colors of error bar and shaded curve). Top: activity in the medial striatum 50-150 ms after stimuli, divided by stimulus side and presence (from **a**). Middle two panels: activity in the centromedial striatum −50-50 ms (upper middle) and in the centrolateral striatum 50-150 ms (lower middle) after movement, divided by movement direction. Bottom panel: activity in the dorsolateral striatum 50-150 ms after outcome, divided by outcome type. Green dots and error bars show measured data. Shaded curves indicate prediction of striatal activity from cortical activity (x-axis values) plus an offset dependent on task event type (stimulus condition, movement direction, rewarded or not). Error bars represent SEM across recordings. * Significant difference across contexts compared to shuffled contexts, p < 0.05, two-tailed. **c**, Mean stimulus responses in trained mice (top row) and naïve mice (bottom row) during passive viewing of task stimuli. Left: mean time course of dorsomedial striatal responses to 100% contrast contralateral (red) and ipsilateral (blue) stimuli. Middle, mean time course of cortical responses to the same stimuli. Right, striatal activity (y-axis) as a function of associated cortical activity (x-axis) and stimulus side (as in **b**, top panel).

This analysis revealed that the relationship between cortical and striatal activity was not fixed, but rather depended on task events specific to each striatal domain (**Fig. 4b**). Activity in the dorsomedial striatal domain was correlated with activity in the corresponding cortical regions regardless of a contralateral, ipsilateral, or invisible stimulus, indicating a consistent component of the corticostriatal relationship (**Fig. 4b, top panel**). Importantly however, there was an additional increase in striatal activity relative to predictions from cortical activity specifically during contralateral stimuli, suggesting a stimulus-dependent deviation in the corticostriatal relationship (**Fig. 4b, top panel, red dots vs. blue and black dots**). The stimulus-evoked boost in striatal activity was approximately independent of cortical activity, suggesting that contralateral stimuli induce an additive boost in striatal activity relative to cortical activity (**Fig. 4b, top panel, shading**). A similar event-specific corticostriatal divergence was observed with respect to movements in the dorsocentral striatum, with an additive increase in striatal activity specifically during contralaterally-orienting movements (**Fig. 4b, middle two panels**). The dorsolateral striatum on the other hand did not exhibit a divergence around outcome despite the outcome producing the largest response (**Fig. 4b, bottom panel**), suggesting that stimulus and movement responses in the dorsomedial and dorsocentral striatum were uniquely divergent from cortical activity.

This stimulus-specific difference between striatal activity and cortical predictions was also observed while mice passively viewed stimuli, but only after learning the task (**Fig. 4c**). We presented the same stimuli used in the task passively, with no opportunity to earn reward, either to some of the trained mice (n = 16 sessions across 2 mice), or to a cohort of naïve mice (n = 23 sessions across 5 mice). Just as during the task, the dorsomedial striatum in trained mice had stronger responses to contralateral stimuli than predicted from the cortex, indicating that this stimulus-induced divergence between the cortex and the striatum did not depend on task engagement (**Fig. 4c, top row**). However, this divergence was absent in naïve mice, where contralateral stimulus responses were still present but the relationship between cortical and striatal activity was consistent across stimulus contexts (**Fig. 4c, bottom row**). These results suggest that learning induces a stable modification in the relationship between cortical activity and striatal activity.

In summary, we found that striatal activity during visually-guided behavior largely mirrors patterns of global cortical activity, suggesting that corticostriatal projections stereotypically propagate cortical activity to associated striatal domains. After task learning, however, striatal responses to contralateral stimuli and movements were additively stronger than predicted from associated cortical responses, regardless of task engagement. Potential mechanisms for this stronger activity include subcortical inputs such as those from the thalamus^18^, learning-induced changes in striatal processing, or a heterogeneous drive from specific subsets of corticostriatal cells^13^. Our results support a view of the corticostriatal circuit where activity in striatal domains reflects patterns of activity in associated cortical regions, with learning inducing adjustments in an otherwise consistent relationship between the two structures.

## Acknowledgements

We thank C. Reddy, M. Wells, L. Funnell, and H. Forrest for mouse husbandry and training, R. Raghupathy for histology, and the NVIDIA Corporation for donation of a Titan X GPU. This work was supported by a Newton International Fellowship, EMBO Fellowship (ALTF 1428-2015), and a Human Frontier Science Program Fellowship (LT226/2016-L) to A.J.P, a Human Frontier Science Program Fellowship (LT001071/2015-L) and Marie Skłodowska-Curie fellowship of the E.U. Horizon 2020 (656528) to N.A.S., Wellcome Trust grants 205093, 204915 to K.DH. and M.C, ERC grant 694401 to K.D.H, M.C. holds the GlaxoSmithKline / Fight for Sight Chair in Visual Neuroscience.

## Author Contributions

A.J.P, K.D.H, and M.C. conceived and designed the study. N.A.S. set up systems for widefield imaging and Neuropixels recordings. A.J.P. collected and analyzed the data. A.J.P., K.D.H., and M.C. wrote the manuscript with input from N.A.S.

---

**Extended Data Figure 1.**
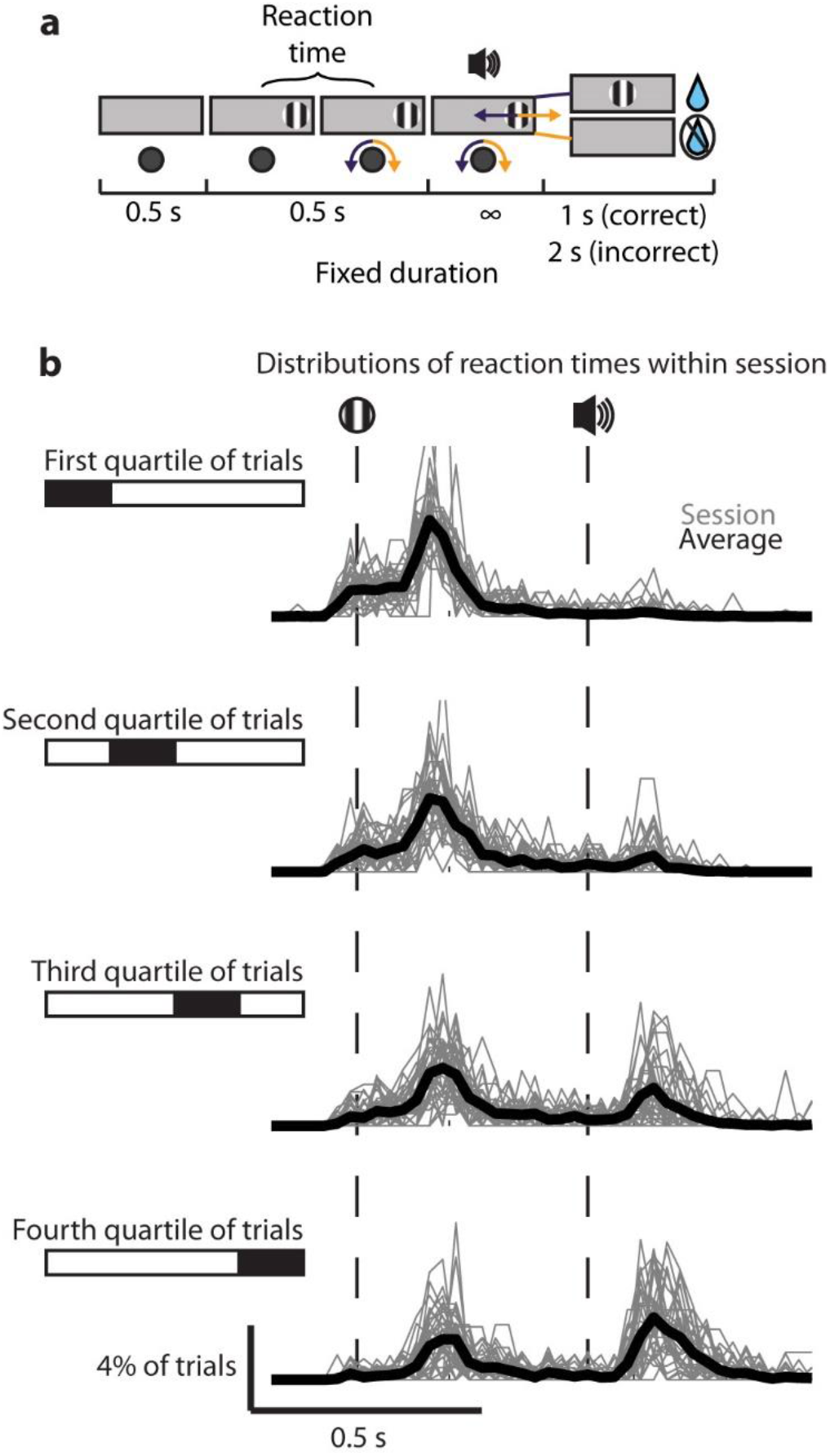
Mice have quick reaction times but sometimes wait for cue late within session. **a**, Timeline of events within trials, **b**, Histogram of times from stimulus to movement onset divided into quartiles of trials per session (top, first 25% of trials within a session; bottom, last 25% of trials within a session). Gray lines denote sessions, black lines denote average across sessions. Dotted vertical lines indicate stimulus onset and go cue tone when stimulus position became yoked to wheel position.

**Extended Data Figure 2.**
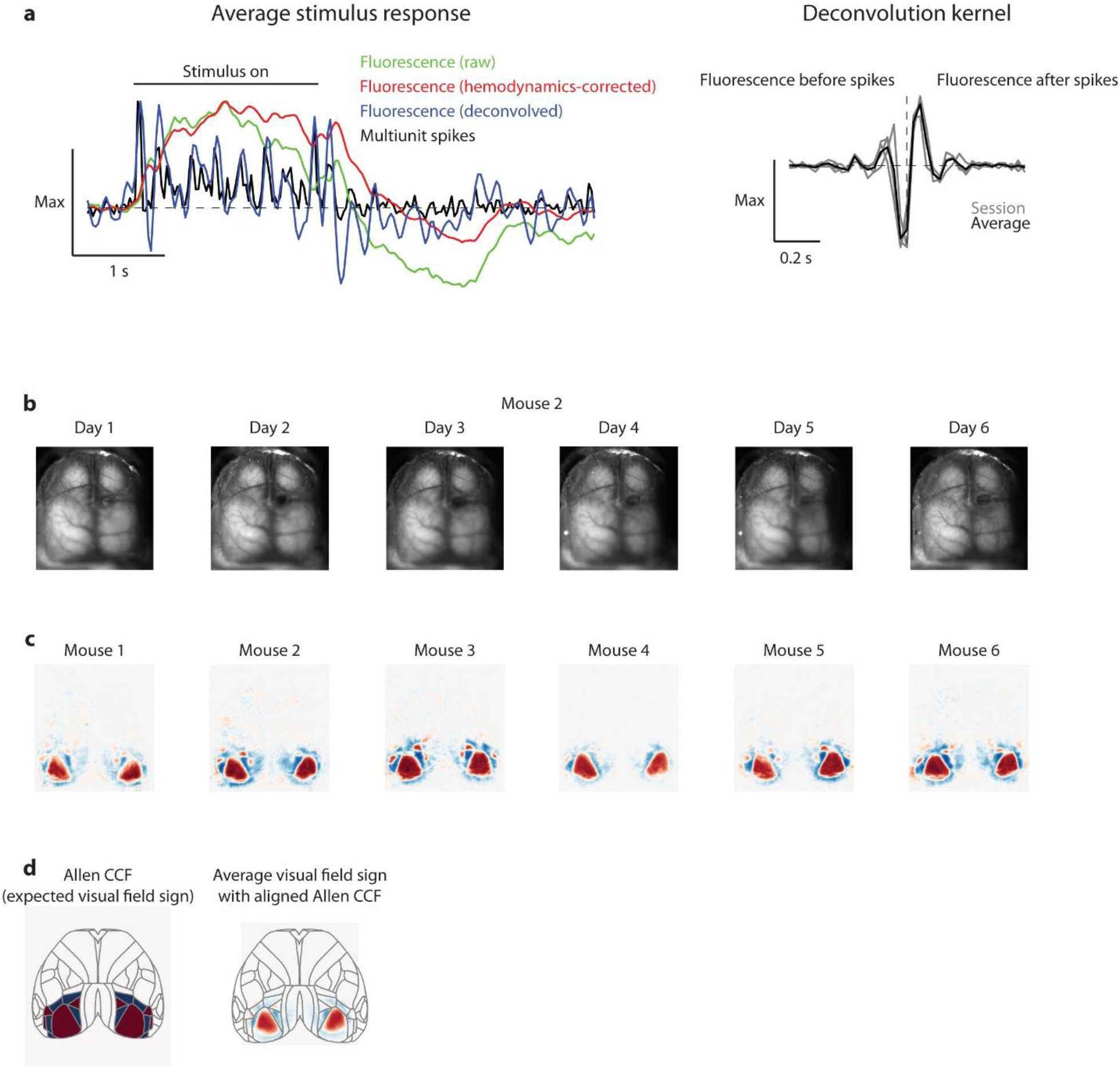
Cortical widefield imaging processing and alignment. **a**, Left, example simultaneous widefîeld fluorescence and multiunit activity within the visual cortex. Traces show average activity aligned to the onset of a large flickering grating presented for 2 s. Black, multiunit activity; green, raw fluorescence (note the slow time course relative to spikes and the large post-stimulus dip); red, hemodynamically-corrected fluorescence (note the reduced post-stimulus dip); blue, deconvolved fluorescence (note the high correlation with spikes). Right, fluorescence-to-spike deconvolution kernels estimated from simultaneous cortical widefield imaging and electrophysiology. Gray, mouse; black, average used for subsequent deconvolution (e.g. used to convert the red line to the blue line in the left panel), **b-d**, Widefield alignment across days, mice, and to the Allen CCF atlas, **b**, Average images are aligned across days within each animal, **c**, Retinotopic visual field sign maps are aligned and averaged across days for each mouse, then average visual sign field maps are aligned across mice, **d**, The CCF is colored according to expected visual field sign (left) and aligned to the average visual field sign map across mice (right). Note that the CCF alignment is for visualization only and is not used for analysis.

**Extended Data Figure 3.**
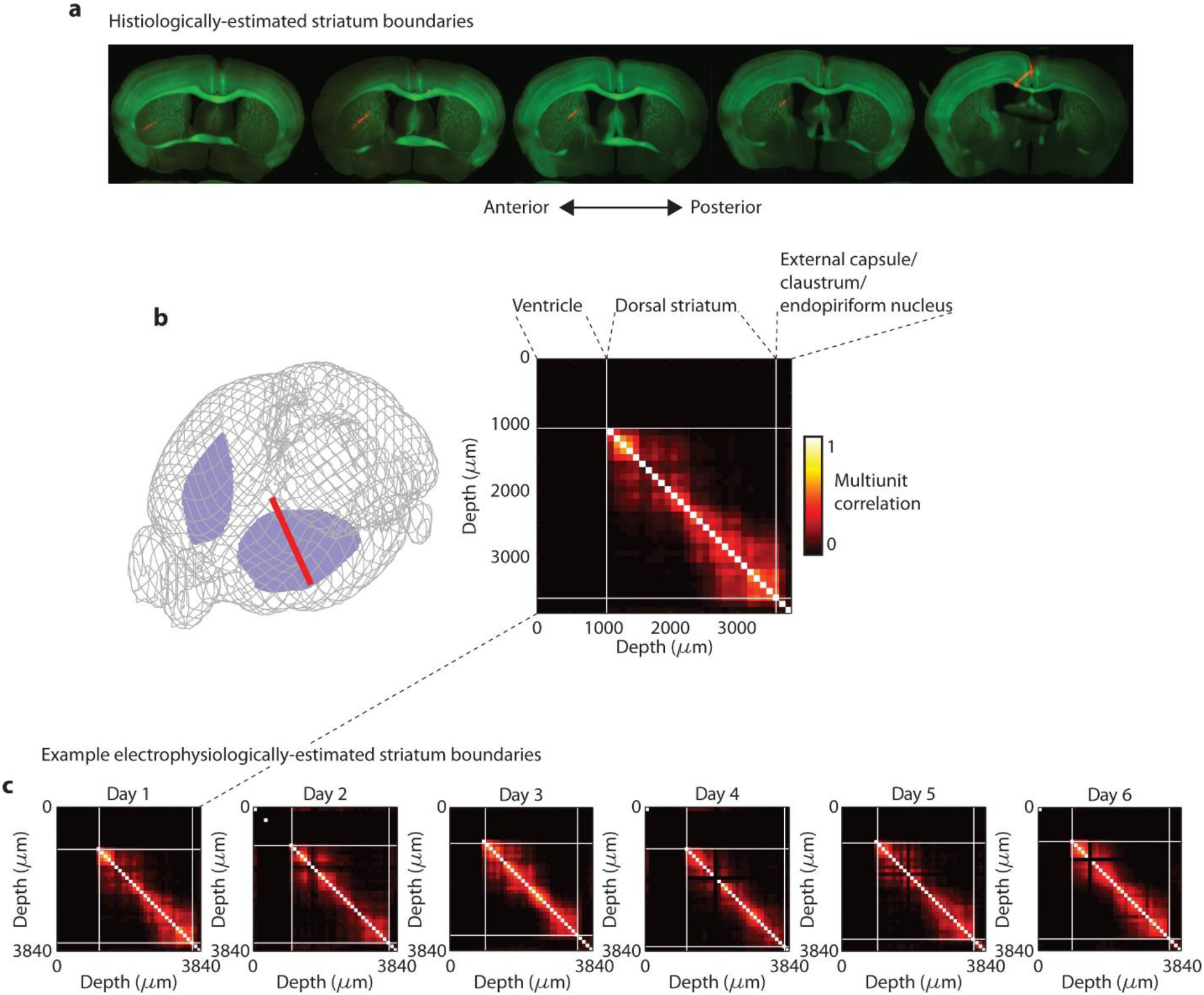
Determining borders of striatum in electrophysiological recordings. **a**, Example histology showing GCaMP6s fluorescence (green) and dye from the probe (red), **b**, Left, reconstructed probe trajectory from (a) in the Allen CCF (left, red: probe, purple: striatum); right, multiunit correlation by depth along the probe from the session when the probe was dyed with brain areas labelled according to histological probe reconstruction, **c**, Example multiunit correlation by depth along the probe for multiple days, with the borders of the striatum approximated medially by the lack of spikes in the ventricle and laterally by the sudden drop in local multiunit correlation.

**Extended Data Figure 4.**
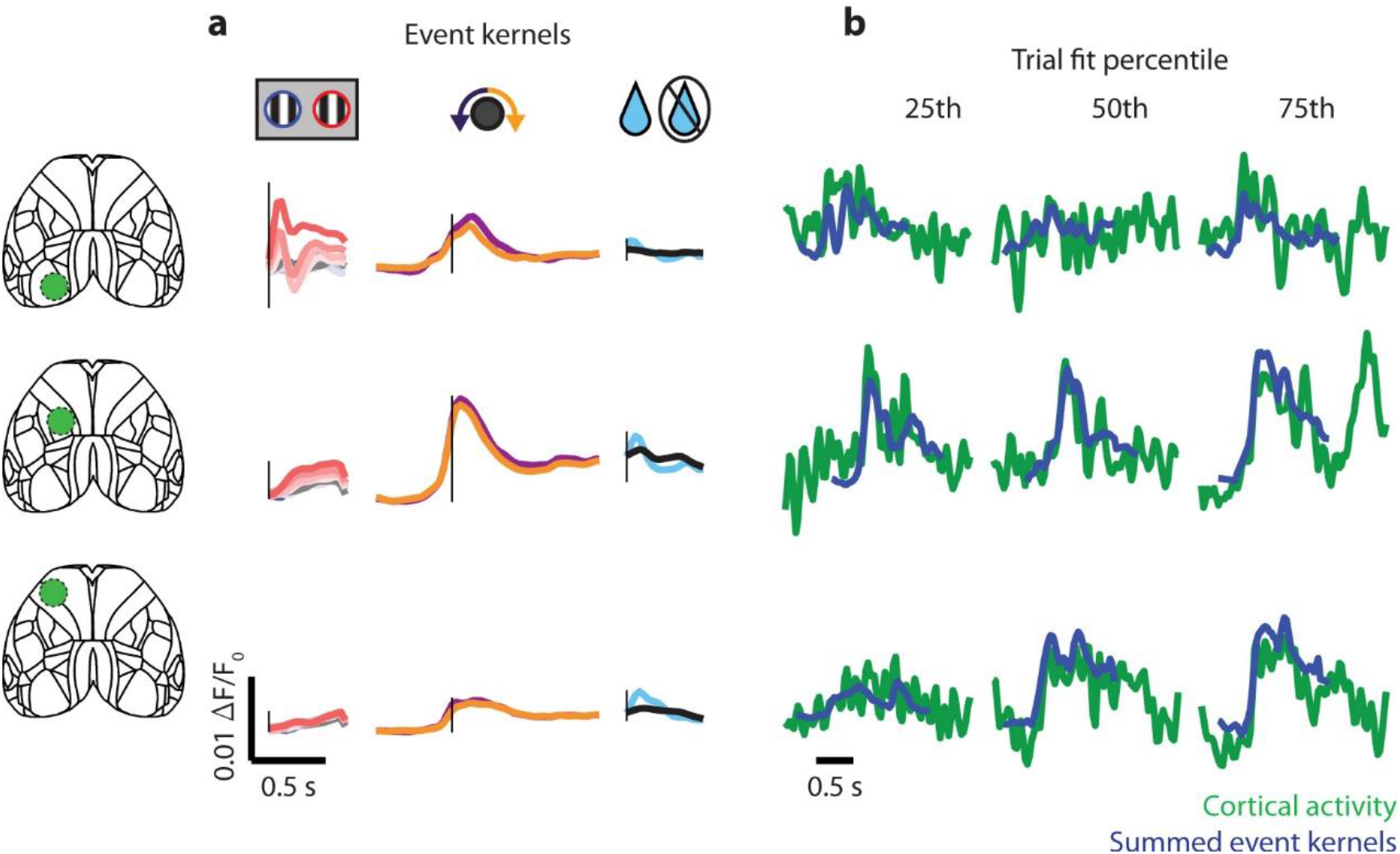
Cortical activity exhibits a sensorimotor gradient of activity. **a**, Kernels from regressing task events to cortical activity for stimuli (left), movements (middle) and outcome (right) averaged within the ROIs indicated on the left, **b**, Example trials of cortical activity (green) and its prediction from task events (blue) within the ROIs as in (a); each column is an example chosen by R^2^ fit percentile. Green traces without overlying blue traces indicate times when there were no overlapping task kernels.

**Extended Data Figure 5.**
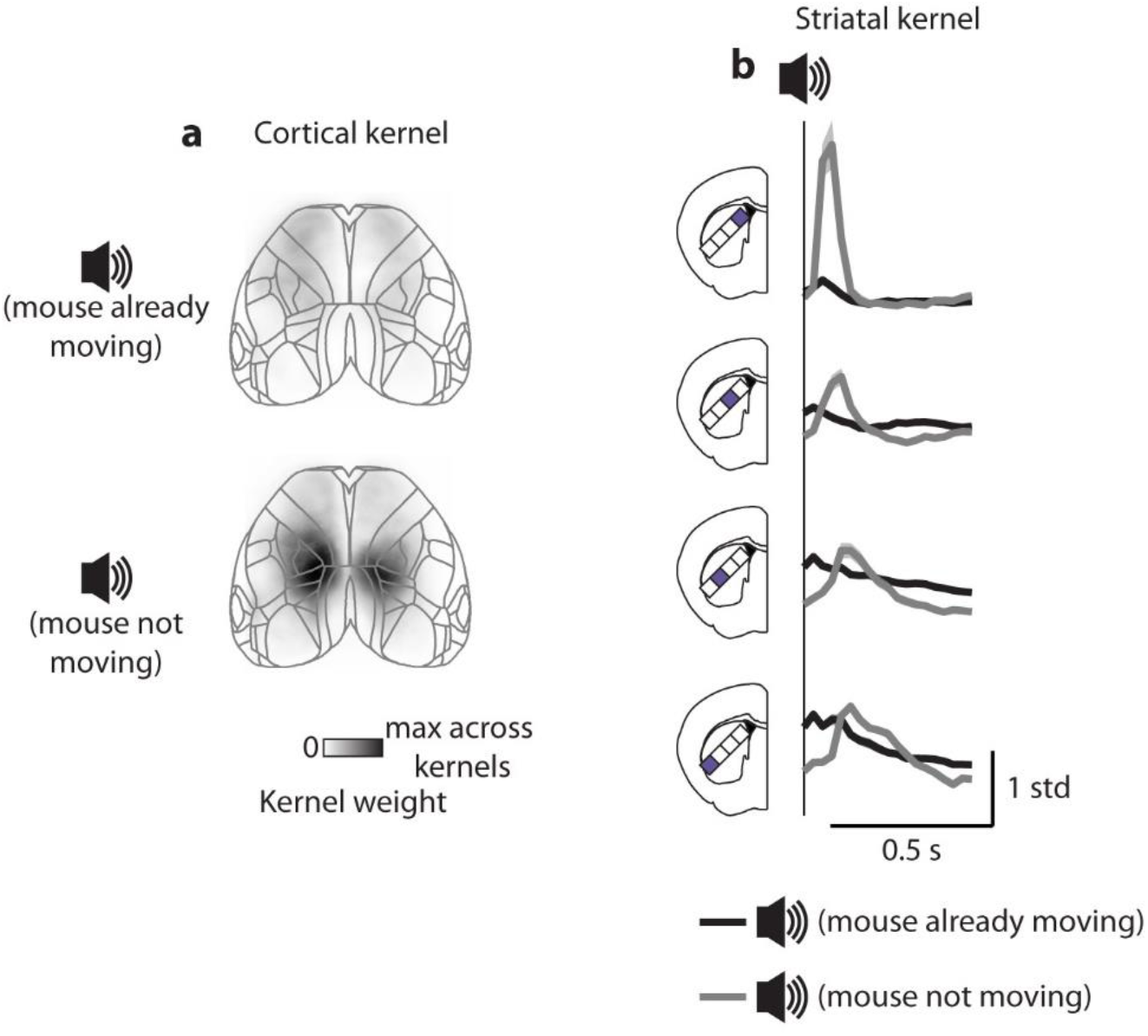
Responses to auditory cue are only on trials with no prior movement. **a**, Example frame from the spatiotemporal kernel in cortical activity (as in Fig. 1d) 50 ms after the go cue tone. Top, accompanying reaction times less than 500 ms (before go cue); bottom, accompanying reaction times greater than 500 ms (after go cue), **b**, Kernel for the go cue in striatal domain activity (as in Fig. 3b) accompanying reaction times less than 500 ms (before go cue, black) or greater than 500 ms (after go cue, gray).

**Extended Data Figure 6.**
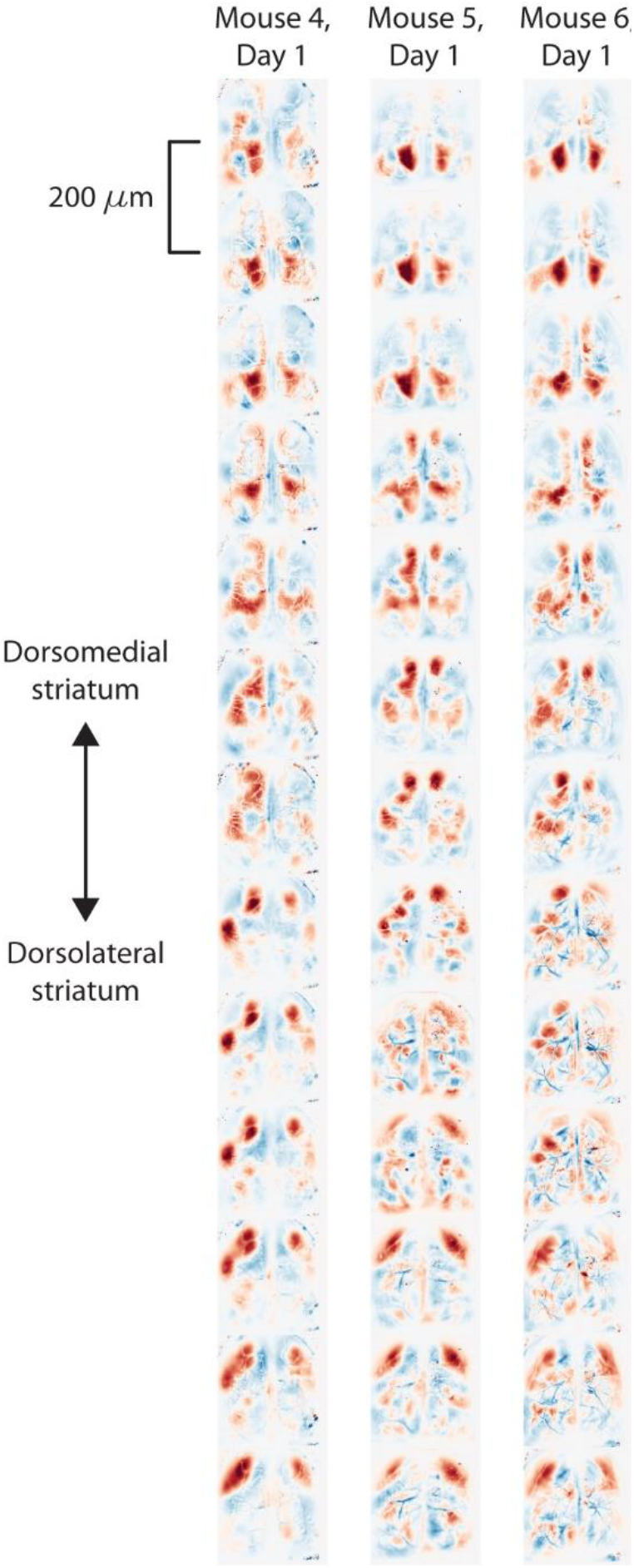
Corticostriatal maps are similar across experiments. Example kernels from regressing cortical activity to striatal activity obtained by successive 200 μm segments along the Neuropixels electrode in one recording session from each of three mice. Spatial map corresponding to spatiotemporal kernel at lag of 0 s is shown, with weights normalized to the standard deviation within each map.

**Extended Data Figure 7.**
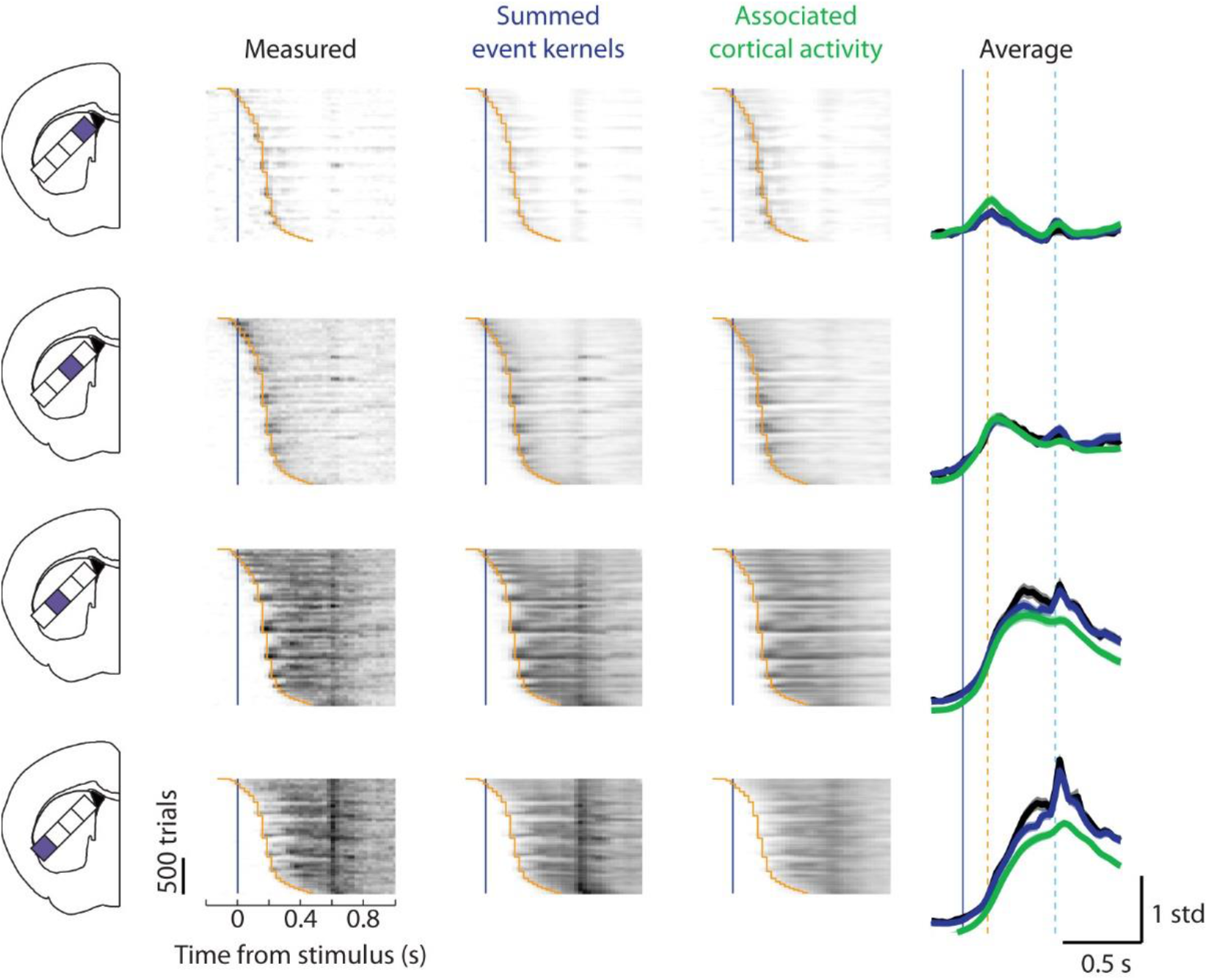
Measured, task-predicted, and cortex-predicted striatal activity during correct ipsilateral trials. Striatal activity for each of the four domains, from measured data (left, black box), predicted from task events (middle, blue box), and predicted from cortical activity (right, green box), as in Fig. 3 but for trials with ipsilateral stimuli and ipsilaterally-orienting movements.

**Extended Data Figure 8.**
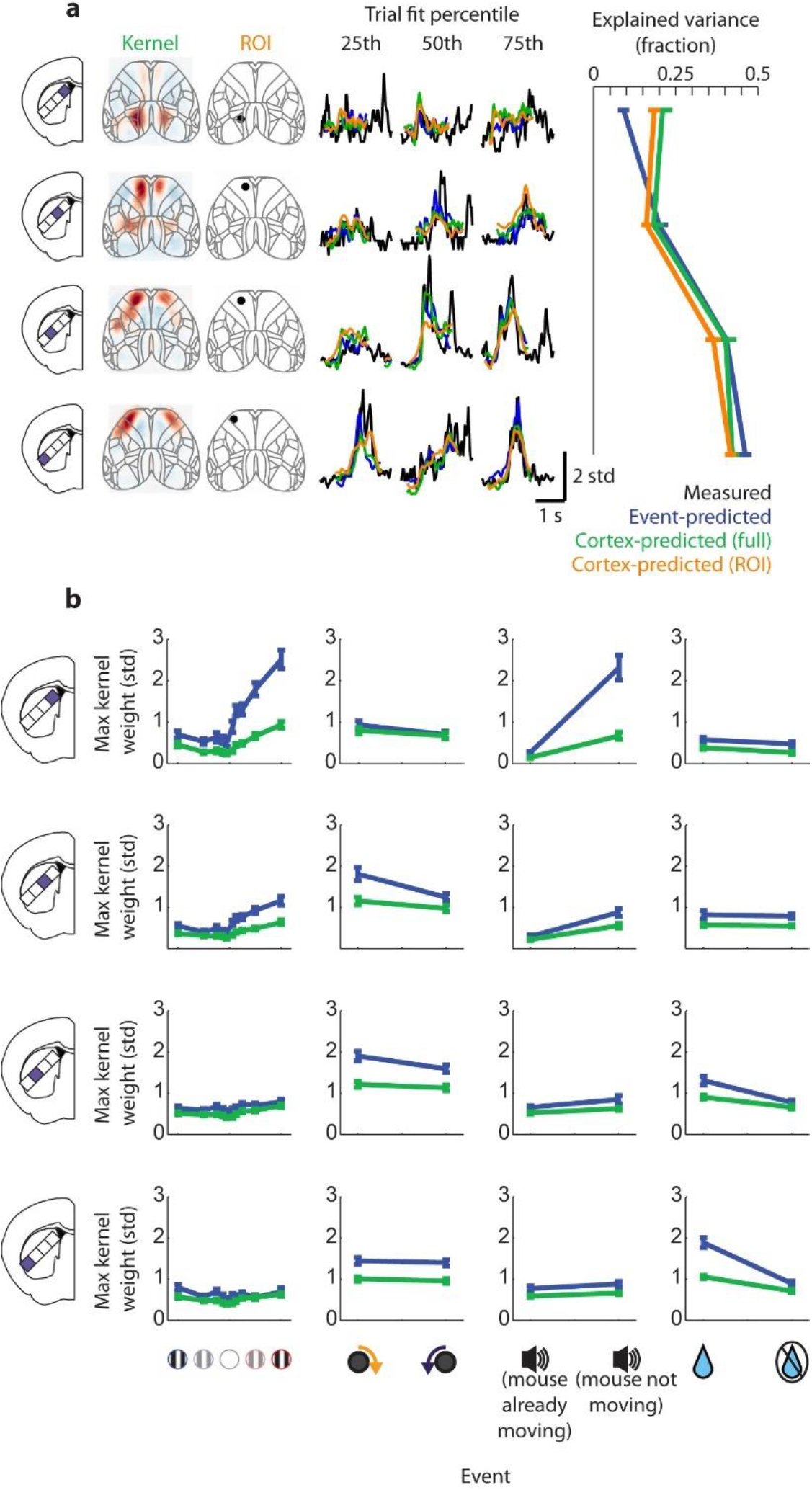
Total and event-related striatal activity explained by task and cortex. **a**, Left, example trials of striatal activity (black), its prediction from task events (blue), from cortical activity using the kernel (green), and from cortical activity within a small region of interest (orange), smoothed with a 100 ms running average filter. Each row represents a striatal domain; each column an example chosen by R^2^ fit percentile from task events. Right, total R^2^ explained variance for prediction from task events (blue), cortical activity using the kernel (green), and cortical activity within a small region of interest (orange) for each striatal domain (y-axis). Cortical kernels accounted for more striatal variance than regions of interest in the first three domains but not in the fourth (paired sign rank test, p = 0.002, 0.02, 0.02, 0.41) **b**, Kernel amplitude (maximum across time) for regressing striatal activity from task events (blue) and domain-associated cortical activity (green). Each row represents a striatal domain, each column an event type.

**Movie 1 | Average cortical fluorescence by trial type.** Fluorescence averaged across recording sessions, color axis from 0-0.02 ΔF/F_0_.

**Movie 2 | Stimulus kernels for cortical fluorescence.** Color normalized to the maximum weight across stimulus kernels.

**Movie 3 | Movement kernels for cortical fluorescence.** Color normalized to the maximum weight across movement kernels.

**Movie 4 | Go cue kernels for cortical fluorescence.** Color normalized to the maximum weight across go cue kernels.

**Movie 5 | Outcome kernels for cortical fluorescence.** Color normalized to the maximum weight across outcome kernels.

**Movie 6 | Preferred cortical activity for striatal domains.** Calculated as the kernel from regressing striatal multiunit activity within each domain from cortical fluorescence. Color normalized to the maximum absolute weight for each kernel.

## Methods

All experiments were conducted according to the UK Animals (Scientific Procedures) Act 1986 under personal and project licenses issued by the Home Office.

### Animals

Mice were adult (6 weeks or older) male and female transgenic mice (TetO-G6s;Camk2a-tTa^33^) which did not show evidence of epileptiform activity^37^.

### Surgery

Two surgeries were performed for each animal, the first being headplate implantation and widefield imaging preparation, and the second being a craniotomy for acute electrophysiology. Mice were anesthetized with isoflurane, injected subcutaneously with Carprieve, and placed in a stereotax on a heat pad. The head was then shaved, the scalp cleaned with iodine and alcohol, and the scalp was removed to expose the skull. The cut skin was sealed with (VetBond, World Precision Instruments), the skull was scraped clean and a custom headplate was fixed to the interparietal bone with dental cement (Super-Bond C&B). A plastic 3D-printed U-shaped well was then cemented to enclose the edges of the exposed skull. A thin layer of VetBond was applied to the skull followed by two layers of UV-curing optical glue (Norland Optical Adhesives #81, Norland Products). Carprieve was added to the drinking water for 3 days after surgery. Mice in the trained cohort were then trained in the task, and after training (or rig acclimation for the naïve cohort) mice were anesthetized and a small craniotomy was drilled over approximately 200 μm anterior and 1000 μm lateral to Bregma.

### Task

Mice were trained on a 2-alternative forced choice task requiring directional forelimb movements to visual stimuli (**Extended Data Fig. 1a**). Mice were headfixed and rested their body and hindpaws on a stable platform and rested their forepaws on a wheel that was rotatable to the left and right. Trials began with 0.5 s of enforced quiescence, where any wheel movements reset the time. A static vertical grating stimulus then appeared 90° from center with a gaussian window σ = 20°, spatial frequency 1/15 cycles/degree, and grating phase randomly selected on each trial. After 0.5 s from stimulus onset, a go cue tone (12 kHz, 100 ms) sounded and the position of the stimulus became yoked to the wheel position (e.g. leftward turns moved the stimulus leftward). Mice usually began turning the wheel before the go cue event on trials with 0% contrast (invisible) stimuli, indicating a rapid decision process and expected stimulus time, although as the session progressed and mice became sated they began waiting for the go cue more often (**Extended Data Fig. 1b**). Bringing the stimulus to the center (correct response) locked the stimulus in the center for 1 s and 2 μL of water was delivered from a water spout near the mouse’s mouth, after which the stimulus disappeared and the trial ended. Alternately, moving the stimulus 90° outward (incorrect response) locked the stimulus in place off-screen and a low burst of white noise played for 2 s, after which the trial ended. The stimulus contrast varied across trials including 0%, 6%, 12.5%, 25%, 50%, and 100%. Difficulty was modulated with an alternating staircase design, where even trials used a random contrast, and odd trials followed a staircase that moved to a lower contrast after 3 correct responses and moved to higher contrast after 1 incorrect response. Correct responses on high-contrast trials were encouraged by immediately repeating all incorrect trials with 50% or 100% contrast, but these repeated trials were excluded from all analyses. Other than repeat trials, stimulus side was selected randomly on each trial. Mice were trained in stages, where first they were trained to ~70% performance with only 100% contrast trials, then lower contrasts were progressively and automatically added as performance increased. Imaging sessions began after all contrasts had been added, and simultaneous imaging and electrophysiology sessions began after ~4 days of imaging-only sessions. Sessions where mice performed less than 85% correct on both left and right 50-100% contrast stimuli were excluded, which resulted in 11 excluded sessions, including all 6 sessions in one mouse which was therefore excluded from analysis.

### Widefield imaging

Widefield imaging was conducted with a sCMOS camera (PCO Edge 5.5) affixed to a macroscope (Scimedia THT-FLSP) with a 1.0x condenser lens and 0.63x objective lens (Leica). Images were collected with Camware 4 (PCO) and binned in 2×2 blocks giving a spatial resolution of 20.6 μm/pixel at 70 Hz. Illumination was generated using a Cairn OptoLED with alternating blue (470 nm, excitation filter ET470/40x) and violet (405 nm, excitation filter ET405/20x) light to capture GCaMP calcium-dependent fluorescence and calcium-invariant hemodynamic occlusion respectively at 35 Hz per light source. Illumination and camera exposure was triggered externally (PCIe-6323, National Instruments) to be on for 6.5 ms including a 1 ms illumination ramp up and down time to reduce light-induced artifacts on the Neuropixels probe. Excitation light was sent through the objective with a 3mm core liquid light guide and dichroic (387/11 single-band bandpass) and emitted light was filtered (525/50-55) before the camera.

Widefield data was compressed using singular value decomposition (SVD) of the form **F** = **USV^T^**. The input to the SVD algorithm was **F**, the *pixels × time* matrix of fluorescence values input to the SVD algorithm; the outputs were **U**, the *pixels × components* matrix of template images; **V** the *time × components* matrix of component time courses; and **S** the diagonal matrix of singular values. The top 2000 components were retained, and all orthogonally-invariant operations (such as deconvolution, event-triggered averaging and ridge regression to predict striatal activity from the widefield signal) were carried out directly on the matrix **V**, allowing a substantial saving of time and memory.

Hemodynamic effects on fluorescence were removed by regressing out the calcium-independent signal obtained with violet illumination from the calcium-dependent signal obtained with blue illumination. To do this, both signals were bandpass Altered in the range 7-13 Hz (heartbeat frequency, expected to have the largest hemodynamic effect), downsampling the spatial components 3-fold, and reconstructing the fluorescence for each downsampled pixel. Pixel traces for blue illumination were then temporally resampled to be concurrent with violet illumination (since colors were alternated), and a scaling factor was fit across colors for each pixel. The scaled violet traces were then subtracted from the blue traces.

To correct for slow drift, hemodynamic-corrected fluorescence was then linearly detrended, high-pass filtered over 0.01 Hz, and ΔF/F_0_ normalized by dividing by the average fluorescence at each pixel softened by adding the median average fluorescence across pixels.

Normalized and hemodynamic-corrected fluorescence was deconvolved using a kernel designed to approximate population spiking activity from widefield GCaMP6s fluorescence. This kernel was estimated using data from separate experiments in which widefield imaging was performed simultaneously with Neuropixels recordings in the visual cortex (**Extended Data Figure 2a;** n = 4 experiments across 4 mice). A kernel optimally predicting cortical multiunit activity from the local widefield fluorescence was estimated by regression, and the kernels resulting from each experiment were averaged to yield a final kernel for deconvolution. The resulting kernel was biphasic (**Extended Data Figure 2a, right panel**), indicating that the deconvolution operation was similar to taking the derivative of the fluorescence trace. Nevertheless, the operations were not identical. For example, deconvolution was able to reproduce sustained responses while the derivative could not.

Widefield images across days for each mouse were aligned by affine registration of each day’s average violet-illumination image which was dominated by vasculature (**Extended Data Fig. 2b**). Widefield images across mice were aligned by affine alignment of average visual field sign maps for each mouse (**Extended Data Fig. 2c**). The Allen Common Coordinate Framework (CCF v.3, © Allen Institute for Brain Science) atlas was aligned to the grand average sign map across mice by assigning expected visual field sign to visual areas^38^ and affine aligning the annotated CCF to the average sign map (**Extended Data Fig 2d**).

When combining widefield data across experiments, data was recast from experiment-specific SVD components into a master cross-experiment SVD component set. These master SVD components were created by aligning and concatenating components **U** from the last imaging-only session of all animals (i.e. no simultaneous neuropixels probe recording), performing an SVD on that concatenated matrix, and retaining the top 2000 components to serve as the master SVD component set. Temporal components (**S * V**) for each experiment were recast by

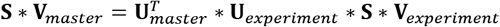

### Electrophysiology

Electrophysiological recordings were made with Neuropixels Phase 3A probes^34^ affixed to metal rods and moved with micromanipulators (Sensapex). Probes were inserted at approximately 200 μm anterior and 1000 μm lateral to bregma at a 45° angle from horizontal (diagonally downwards) and 90° from the anterior-posterior axis (straight coronally) to a depth of about 6 mm from the cortical surface to reach the contralateral striatum. Electrophysiological data was recorded with Open Ephys^39^.

Raw data within the action potential band (soft high-pass filtered over 300 Hz) was de-noised by common mode rejection (i.e. subtracting the median across all channels), and spike-sorted using Kilosort v2 (www.github.com/MouseLand/Kilosort2). Units representing noise were manually removed using phy^40^.

The borders of the striatum were identified within each recording using the ventricle and dorsolaterally-neighboring structure (likely the endopiriform nucleus) as electrophysiological landmarks. Since no units were detected in the ventricle, the start of the striatum on the probe was marked as the first unit after at least a 200 μm gap from the last unit (or the top of the probe if no cortical units were detected). Detected units were continuous after the ventricle, but multiunit correlation across depths of the probe in temporal bins of 10 ms and spatial bins of ~ 100 μm revealed a sharp border in correlation at a location consistent with the end of the striatum (**Extended Data Fig. 3c**). This border was present in every recording and used to define the end of the striatum on the probe.

Probe trajectory was reconstructed (**Extended Data Fig. 3b**) using the GUI from (ref.^41^).

Electrophysiological recordings were synchronized to widefield data and task events by aligning to a common digital signal randomly flipping between high and low states (produced from an Arduino) accounting for both clock offset and drift.

### Regression from task events to activity

Regression from task events to striatal multiunit activity or deconvolved cortical fluorescence activity using linear regression of the form

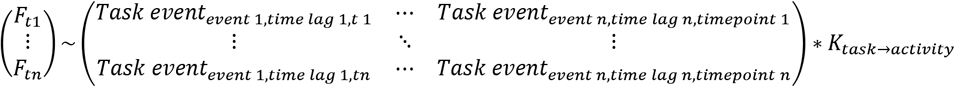

Here, *K_task→actlvlty_* represents a vector containing the concatenated estimated kernels for each event type, estimated by least squares using MATLAB’s \ operator. *F*_*t*1_ to *F*_*tn*_ represent the fluorescence or firing rate time course to be predicted, “baseline-subtracted” by subtracting the average activity 0.5-0 s before stimulus onset, during which time the animals were required not to turn the wheel. For each event type, a task matrix was constructed as a sparse Toeplitz matrix with a diagonal series of Is for each event at each time lag, with zeros elsewhere. Toeplitz matrices were made for each event type: stimulus onset (one for each stimulus side*contrast, lags of 0-0.5 s), movement onset (one each for left and right final response, lags of −0.5-1 s), go cue onset (one for trials where mice had already begun moving and one for trials with no movement before the go cue, lags of 0-0.5 s), and outcome (one for water and one for white noise, lags of 0-0.5 s). These matrices were horizontally concatenated to produce the matrix shown in the above equation. Regression was 5-fold cross-validated by splitting up timepoints into consecutive chunks.

### Regression from cortical activity to striatal activity

Normalized, hemodynamically-corrected, and deconvolved widefield fluorescence was regressed to cortical multiunit activity using ridge regression. Regression took the form

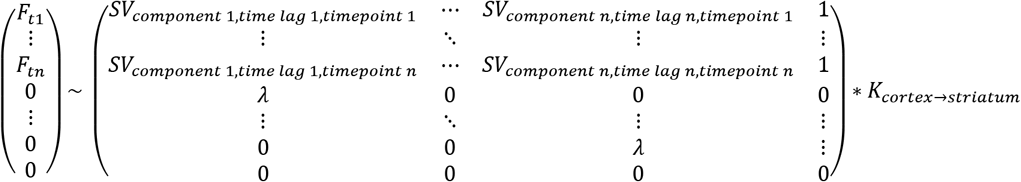

Here, *F*_*t*1_ to *F_tn_* represent the standard-deviation-normalized striatal spiking time course to be predicted. K_corte→striatum_ represents the estimated spatiotemporal kernel from cortical fluorescence to standard-deviation-normalized striatal spiking estimated by least squares using MATLAB’s \ operator. To make the design matrix, a Topelitz matrix was constructed for each temporal SVD component of the cortical widefield, scaled by the singular values (S*V), staggered across a range of time values (−500 ms to +500 ms). These Toeplitz matrices were horizontally concatenated, also including a column of ones to allow an offset term. To regularize using ridge regression, this matrix was vertically concatenated above diagonal matrix of regularization values *λ*, and the striatal activity time courses *F* were concatenated above the same number of zeros. Regression was 5-fold cross-validated by splitting up timepoints into consecutive chunks, and values for *λ* were determined empirically for each experiment by regressing from cortical fluorescence to multiunit from the whole striatum across a range of *λ* values and finding the *λ* that yielded the largest cross-validated explained variance.

### Striatal spike grouping by domain

Striatal domains were defined from their cortical kernels by regressing cortical fluorescence to striatal multiunit as described above, for consecutive 200 μm segments of the Neuropixels track through the striatum recorded in each experiment. The spatial kernels at a time lag of 0 where then combined across all experiments and split into 4 groups through K-means, and the average spatial map for the 4 groups was used as a template for a striatal domain. The number of groups was set at 4 because more groups produced inconsistent K-means results and produced multiple groups of striatal activity with similar activity patterns, while fewer groups did not accurately reflect the diversity of observed maps or activity patterns. The spatial map from each 200 μm striatal segment was then assigned as one of the four groups by highest correlation with the template maps. In order to ensure smooth domain transitions which followed a consistent order by depth (1 being most medial and 4 being most lateral), The domain assignments were median filtered in 3-segment windows which prevented rapid fluctuations and then any assignment which represented a backwards step (e.g. 1 appearing after 2) was replaced with its nearest neighbor assignment.

### Allen connectivity maps

Anatomical projections were labeled using the Allen connectivity database^36^(**Fig. 2c,d**). This was done by defining the targeted trajectory within the striatum in the Allen CCF, splitting that striatal trajectory into 4 equal parts, and querying the Allen API (2015) for injection sites within the cortex that yielded axon terminals at the center of each striatal segment. To maximize coverage across the brain since the Allen connectivity database has different left and right hemisphere injections, queries were performed for striatal sites bilaterally and results from the right striatum were mirrored and combined with the results from the left striatum. Cortical sites with striatal projections in each segment were plotted as dots with size scaled by projection density as returned by the API.

